# Cellular heterogeneity in DNA alkylation repair as a trade-off between cell survival and genetic plasticity

**DOI:** 10.1101/2021.05.24.445533

**Authors:** Maxence S. Vincent, Stephan Uphoff

## Abstract

DNA repair mechanisms fulfil a dual role, as they are essential for cell survival and genome maintenance. Here, we studied how cells regulate the interplay between DNA repair and mutation. We focused on the *Escherichia coli* adaptive response that increases resistance to DNA alkylation damage. Combination of single-molecule imaging and microfluidic-based single-cell microscopy showed that noise in the gene activation timing of the master regulator Ada is accurately propagated to generate a distinct subpopulation of cells in which all proteins of the adaptive response are absent. Although lack of these proteins causes extreme sensitivity to alkylation stress, cellular heterogeneity in DNA alkylation repair provides a functional benefit by increasing the evolvability of the whole population. We demonstrated this by monitoring the dynamics of nascent mutations during alkylation stress as well as the frequency of fixed mutations that are generated by the distinct subpopulations of the adaptive response. This highlighted that evolvability is a trade-off between mutability and cell survival. Stochastic modulation of DNA repair capacity by the adaptive response solves this trade-off through the generation of a viable hypermutable subpopulation of cells that acts as a source of genetic diversity in a clonal population.

## Introduction

Genome plasticity is essential for adaptation of cells to new environments. For instance, bacteria rely on mutagenesis to evolve resistance to antibiotics [1–3] and to adapt to new host environments [4]. On the other hand, maintenance of genome stability is also necessary for their survival. Hence, cells employ conserved genetic networks and stress responses to regulate repair of their DNA [5]. Perturbation of DNA repair pathways by mutations or drug treatments increases the mortality and mutation rates of cells in the presence of DNA damage. Loss of repair functionality can have beneficial consequences for bacterial populations, as an increased mutation rate can enhance evolvability. Indeed, mutator strains consistently evolve during laboratory evolution experiments [6,7], and are frequently found in bacterial isolates from infected patients or the environment [8,9]. These phenotypes have been shown to arise from mutations in DNA mismatch repair, oxidative DNA damage repair, and DNA replication proofreading genes. However, although an increased mutation supply can accelerate adaptive evolution when a population is maladapted in its current environment, inactivation of genome maintenance mechanisms can lower cell fitness and lead to accumulation of deleterious mutations [10]. Besides the existence of permanent genetic mutator alleles, growing evidence suggests that cells can adopt transient hypermutable phenotypes by regulating the expression or activity of DNA repair enzymes [11–14]. Temporary upregulation of mutagenesis is believed to promote evolutionary adaptation in response to stress without compromising genetic stability in optimal environments [15,16]. Furthermore, cell subpopulations with elevated mutation rates could serve as reservoirs of increased genetic plasticity. Despite the compelling logic of this theory, whether a hypermutable subpopulation contributes significantly to the overall evolvability of the whole population depends not only on its mutation rate but also on its size, lifetime, and viability [17]. These crucial parameters are not accessible from conventional genetics assays. As such, it remains unclear if transient hypermutable phenotypes can provide evolutionary benefits, and how any such benefits compare to the evolvability of permanent genetic mutator strains.

Among the broad class of damaging compounds that can generate mutagenic DNA lesions, alkylating agents are found in the external environment [18] and are endogenously produced [19,20]. They can alter nucleobases and phosphotriester linkages of ssDNA, dsDNA and RNA in eukaryotic and prokaryotic cells [21–24]. In *E. coli* six genes have been identified to protect DNA specifically against alkylation damage. The two constitutively expressed enzymes Ogt (O6meG methyltransferase) and Tag (3meA DNA glycosylase I) provide a basal repair capacity [25–28], whereas the four adaptive response components, Ada (O6meG methyltransferase), AlkA (3meA DNA glycosylase II), AlkB (3meC dioxygenase) and AidB are induced upon alkylating stress [29–34]. *ada* and *alkB* are expressed in an operon, while *alkA* and *aidB* have separate promoters (**Fig.1 A**).

**Fig.1:**
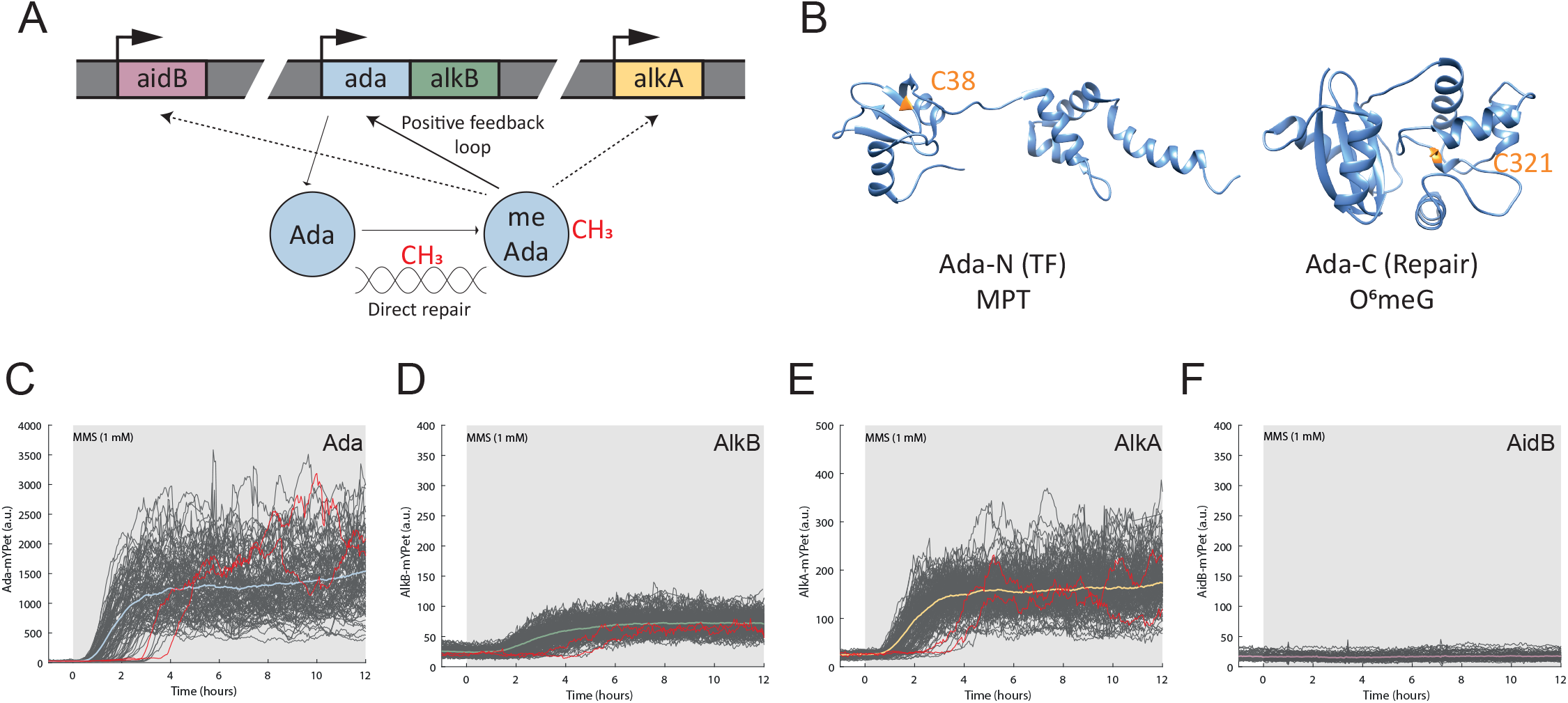
Stochastic activation of Ada affects the expression of the adaptive response genes. (A) Schematic of the adaptive response regulation. The adaptive response gene network is composed of the *ada-alkB* operon, *alkA* and *aidB*. Methylation of the damage sensor protein Ada turns itself into a transcriptional activator the regulon. (B) Ada N-terminal domain (PDB: 1ZGW) and C-terminal domain (PDB: 1SFE) carry the methylated phosphotriester (MPT) and O^6^meG repair activities respectively. The methyl acceptors C38 and C321 are shown in orange. (C-F) Microfluidic-based imaging of the adaptive response components activation. Single-cell time-traces of Ada-mYPet (cells = 104) (C), AlkB-mYPet (cells = 265) (D), AlkA-mYPet (cells = 228) (E) and AidB-mYPet (cells = 146) (F) upon 1 mM MMS treatment (shaded background). Example of cells delaying gene expression are shown in red. Colored curves represent cell average time trace.

The adaptive response is regulated through the methylation status of Ada (**Fig.1 A**). Ada is a bifunctional enzyme, exhibiting a transcription factor (TF) activity carried by its N-terminal domain (Ada-N) and an O^6^meG methyltransferase activity with the catalytic cysteine 321 (C321) in the C-terminal domain (Ada-C) (**Fig.1 B**). Ada-N repairs methylated phosphotriester (MPT) lesions by direct and irreversible transfer of the methyl group onto its catalytic cysteine 38 (C38). The methylation of C38 turns Ada into a transcriptional activator of the adaptive response gene network, which includes its own gene and thus leads to amplification of gene expression by positive feedback [18,35,36].

Although the adaptive response has been characterised for decades, recent single-cell measurements uncovered unexpected cell-to-cell heterogeneity in Ada abundance [37]. Specifically, Ada exhibits large variation in gene expression between cells of isogenic *E. coli* populations [37]. As a result of gene expression noise, the basal level of Ada in absence of alkylating stress is heterogeneous, with a subpopulation of cells containing not even a single molecule of Ada. Consequently, upon alkylating stress, cells devoid of Ada are unable to activate the adaptive response until the stochastic expression of at least one Ada molecule, which can take multiple cell generations [37,38]. Cells with a delayed adaptive response exhibit higher rates of DNA replication errors, suggesting that they could act as a hypermutable subpopulation [37,39]. However, as phenotypic variation is ubiquitous in bacteria, it is difficult to know whether the heterogeneity in the adaptive response genuinely represents a beneficial evolutionary strategy or if it is a side-effect of unavoidable molecular noise. Here, we addressed this question by studying the regulation of the adaptive response and its effects on population evolvability.

## Results

### Stochastic activation of Ada propagates across the whole adaptive response regulon

Alkylation stress causes mutagenic DNA lesions, that promote error-prone DNA replication, and toxic lesions, that block DNA replication forks and lead to cell death if left unrepaired [35]. Indeed, *E. coli* strains with deletions of individual genes of the adaptive response, namely *ΔalkA ΔalkB, and Δada-alkB* (lacking the entire *ada-alkB* operon), were unable to grow on plates in the presence of the alkylating agent methyl methanesulfonate (3 mM MMS) that causes both mutagenic and toxic lesions (**Supp.1**) [35]. However, deletion of *aidB* did not affect cell survival (**Supp.1**). Considering the importance of the adaptive response for tolerance of alkylation stress, it is surprising that the master regulator Ada is so sensitive to gene expression noise that its feedback autoregulation generates large variation in Ada abundances across cells in a population after exposure to MMS [37]. As AlkA and AlkB are crucial for survival of alkylation damage (**Supp.1**), we asked whether the stochastic activation of Ada also impacts their expression. In principle, variation in the master regulator could be buffered or propagated in the gene regulatory network. To this end, we monitored the endogenous expression of functional Ada, AlkB, AlkA and AidB translational fusions to the fluorescent protein mYPet (**Supp.1**) at the single-cell level in a microfluidic device. The ‘mother machine’ setup allows imaging hundreds of single cells over tens of generations under constant growth conditions and defined stress treatments [10,39,40]. We observed that the addition of MMS in the fluidic system caused most cells to activate the adaptive response regulon rapidly (termed ON-state, **Fig.1 C-E**). However, we detected a fraction of cells that delayed the activation of AlkB and AlkA expression for the duration of multiple cell cycles (termed OFF-state), despite constant treatment with a fixed concentration of MMS (**Fig.1 D-E**). This cell-to-cell heterogeneity in *alkB* and *alkA* gene induction matches the patterns seen for *ada* (**Fig. 1C-E**) [37]. In the conditions of our experiments, we did not detect any activation of AidB expression in response to MMS (**Fig.1 F**). Indeed, the role of *aidB* in DNA repair has been brought into question before [35,41,42], and its contribution to the alkylating stress response appears to be negligible considering that a *ΔaidB* strain has the same MMS sensitivity as the wild-type (**Supp.1**).

### Fluctuations in *ada* expression are accurately propagated to *alkA*

The similarity of Ada and AlkB activation timing was expected because both genes are in the same operon, however, the variability observed for AlkA activation was less anticipated since both unmethylated and methylated forms of Ada have been proposed to activate AlkA [43,44]. Thus, to precisely quantify the activation times of Ada and AlkA in the same cell, we engineered dual reporter strains expressing endogenous Ada-CFP and AlkA-mYPet fusions (**Fig.2 A-B and Supp. 1**). We observed that both Ada and AlkA expression share highly correlated activation times (**Fig.2 C**). On closer inspection, we detected a ~40min activation delay for AlkA with respect to Ada (**Fig.2 D**), which indicates that Ada needs to rise in concentration first before it activates *alkA* transcription. To confirm this difference in activation times, we monitored AlkA-mYPet and an ectopic transcriptional *P*_*ada*_-CFP fluorescent reporter. In this strain, the endogenous *ada* allele is unaltered and the activation of AlkA and *P*_*ada*_ became almost simultaneous, thereby confirming our hypothesis (**Supp.2 A,B**). We further noted that Ada and AlkA both displayed broad fluctuations in expression level in single cells with constant MMS treatment even when the cell-average expression had reached steady-state after the period of response activation (**Fig.2 B**). We previously showed that the steady-state fluctuations of Ada reflect variation in the amount of DNA damage in individual cells over time [37]. Temporal cross-correlation between Ada and AlkA signals showed that fluctuations of *ada* expression are correlated with those of *alkA* (**Fig.2 F and Supp.2 C**). As a control, we did not detect cross-correlations between Ada and AlkA signals from different random cells or between AlkA and an unrelated P_RNA1_-mKate2 fluorescent reporter (**Fig.2 F and Supp.2 C**).

**Fig.2:**
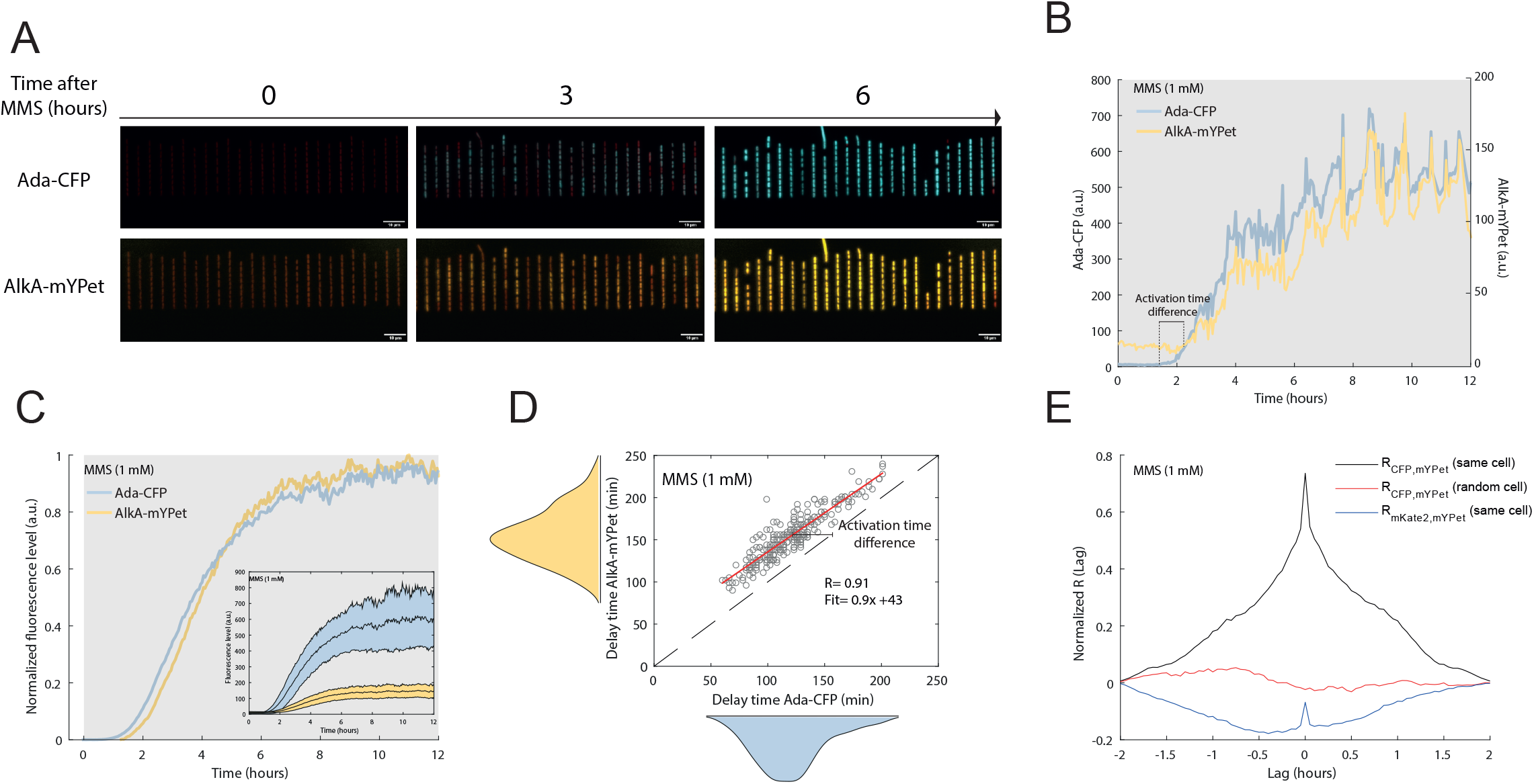
Fluctuations in *ada* expression are propagated to *alkA*. Dual reporter assays of Ada-CFP and Alka-mYPet expressions. (A) Example of microfluidic single-cell imaging of the dual reporter strain carrying the Ada-CFP and AlkA-mYPet reporters with constant 1 mM MMS treatment. Ada-CFP fluorescence is displayed in blue, AlkA-mYPet is displayed in yellow and constitutive mKate2 fluorescent cell marker is displayed in red. (B-C) Example of single-cell time traces showing activation of Ada-CFP (B) and AlkA-mYPet (C) after 1 mM MMS addition (shaded background). (D) Fluorescence of each single-cell (cells = 239) has been averaged, normalized and subtracted from their level at time = 0h (addition of MMS in the microfluidic system). Inset shows the original signals and their standard deviations about the mean (colored regions). (E) Correlation plot showing delay times between 1 mM MMS addition and response activation for Ada-CFP and AlkA-mYPet. Each circle represents one cell. R is the Pearson coefficient. The red line shows the best linear fit. (F) Cross-correlations of Ada-CFP and AlkA-mYPet signals between 9 and 11 hours after MMS addition. The average of each individual cross-correlation between the mYPet and CFP signals from the same cell is represented by the black curve, whereas the red curve represents the average from two random cells and indicates that the correlation between Ada and AlkA is specific of their respective cell. The blue curve represents the average of each individual cross-correlation between the AlkA-mYPet signal and the segmentation marker mKate2 signal from the same cell and indicates that the correlation is independent of the fluorescence fluctuations due to cell elongation during the cell cycle.

### The basal level of the adaptive response proteins is low and heterogeneous

The propagation of stochastic Ada activation to the whole response regulon means that the cells with a delayed response (the OFF subpopulation) dwell in a state in which all proteins of the Ada regulon are only expressed at a basal level. We therefore quantified the basal expression of these proteins using a method to count translational protein fusions to the HaloTag, which can be labelled with the fluorescent ligand TMR [26]. MMS sensitivity assays confirmed the functionality of the HaloTag fusions (**Supp.1**). Chemical fixation of cells allowed us to capture long camera exposures on a custom-built single-molecule fluorescence microscope such that we were able to detect distinct fluorescent spots and count protein copy numbers per cell (**Supp.3**). The basal expression of Ada has been previously shown to be as low as 1 molecule/cell on average [37,45], which was similar to the distribution of Ada-Halo molecules/cell that we observed here (**Fig.3 A**). Single-molecule counting of AlkB-Halo revealed that most cells were completely devoid of AlkB in absence of alkylating stress (**Fig.3 B**). Only ~20 % of the population exhibited a single AlkB protein (**Fig.3 B**). This observation is surprising given the importance of AlkB for the repair of alkylation damage (**Supp.1**), however, it is not unexpected considering that *alkB* is positioned at the end of the *ada-alkB* operon and likely to be less transcribed than *ada*. We further quantified the absolute number of AlkA-Halo proteins (**Fig.3 C**). Although some cells (~5% of the population) contained too many proteins (>8) to be accurately counted, most cells in the population exhibited a low number of AlkA, with ~2.6 molecules per cell on average. As for AlkB, it is surprising that the important DNA repair protein AlkA is expressed at such low levels. Of note, we did not detect any AidB-Halo proteins in most cells (>95% of the population) (**Fig.3 D**). Overall, these results demonstrate that AlkA and AlkB are necessary for the cell to survive alkylation stress (**Supp.1**) yet they are present at very low level and in many cells completely absent before induction.

**Fig.3:**
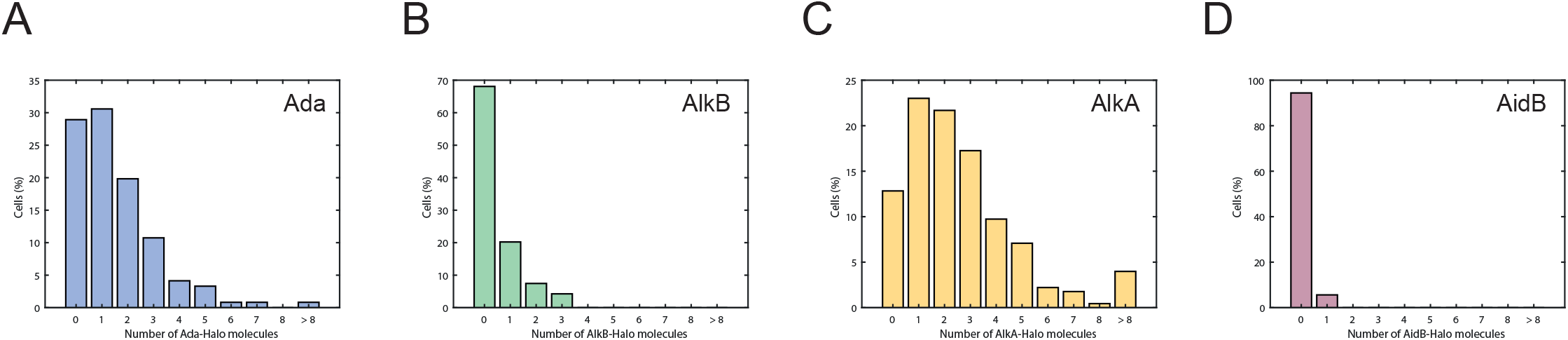
Basal level of the adaptive response proteins. The distribution of Ada-Halo (cells= 121), AlkB-Halo (cells= 94), AlkA-Halo (cells= 238) and AidB-Halo (cells = 105) proteins per cell are shown in panel A, B, C and D respectively.

### Phenotypic heterogeneity of the adaptive response appears to be an evolved property

Despite the apparent noisiness of the adaptive response, our results demonstrate that it is in fact a remarkably precise gene regulatory network that splits an isogenic population of cells into two phenotypically distinct and defined subpopulations. The production of single Ada molecules functions as the stochastic master switch in this network. The random timing of Ada activation in each cell is precisely transmitted to induce AlkB and AlkA expression after a further delay (**Fig.1 and Fig.2**). Cells with delayed Ada activation are essentially devoid of all adaptive response proteins because of their extremely low basal expression levels (**Fig.3**). Therefore, the OFF state is distinct and defined not just by the absence of the Ada regulator, but an all-round lack of proteins that are crucial for DNA alkylation repair. After switching to the ON state, fluctuations in Ada production are propagated such that the whole response regulon (except AidB) closely follows the state of the regulator (**Fig.1 and Fig.2**). These conclusions suggest that the stochastic phenotypic heterogeneity generated by the adaptive response is an evolved property of the system, rather than side-effect of a regulatory inaccuracy.

### Contribution of Ada, AlkB, and AlkA to genome maintenance

The fact that lack of AlkB and AlkA is very toxic to cells in the presence of alkylation stress (**Supp. 1**) suggests that the formation of a distinct subpopulation of cells in which these proteins are absent must have a functional purpose that outweighs the fitness costs. Nevertheless, whether heterogeneity in the adaptive response is exploited as a functional benefit for a cell population remains an open question. An interesting hypothesis is that the delay of Ada, AlkB, and AlkA activation could increase the mutation rate of certain cells and therefore provide an adaptive and heritable genetic diversity (also referred to as evolvability [46]). We previously showed that cells with a delayed adaptive response have a higher rate of DNA replication errors during MMS treatment than cells that rapidly activated the response [37,39]. To address the functional benefits of such a mechanism, we determined the contribution of each component of the adaptive response regulon to genome maintenance under alkylating stress. We used a method that enables the detection of nascent DNA replication errors making use of the fluorescently-labelled MutL-mYPet fusion protein that forms distinct fluorescent foci when bound at DNA mismatches (**Fig.4 A**) [10,39]. As shown before, delayed activation of the adaptive response causes a transient burst in the rate of DNA mismatches that lasts for ~2 hours after the addition of 1 mM MMS [39]. Unlike the wild-type, strains with the gene deletions *ΔalkB, ΔalkA*, and *Δada-alkB* all showed elevated and sustained mismatch rates that did not recover during prolonged MMS treatment (**Fig.4 B**). Addressing the specific function of Ada in DNA repair is more complex than for AlkB and AlkA, because Ada has a dual role as an O^6^meG repair protein and regulator of the adaptive response. To separate these functions, we engineered an endogenous chromosomal Ada mutant, Ada^C321A^, that lacks the catalytic cysteine required for repair of O6meG lesions (**Supp.1**). This mutant is still able to regulate the adaptive response, which is activated by methylation of Cys38 [36,47,48]. Upon alkylating stress, O6meG repair deficiency resulted in a sustained and increased mismatch rate with respect to the WT level, but remained below the *ΔalkB* and *ΔalkA* levels (**Fig.4 B**). Therefore, AlkB, AlkA, and Ada each provide specific DNA repair functions that are important for mutation prevention during alkylation stress.

**Fig.4:**
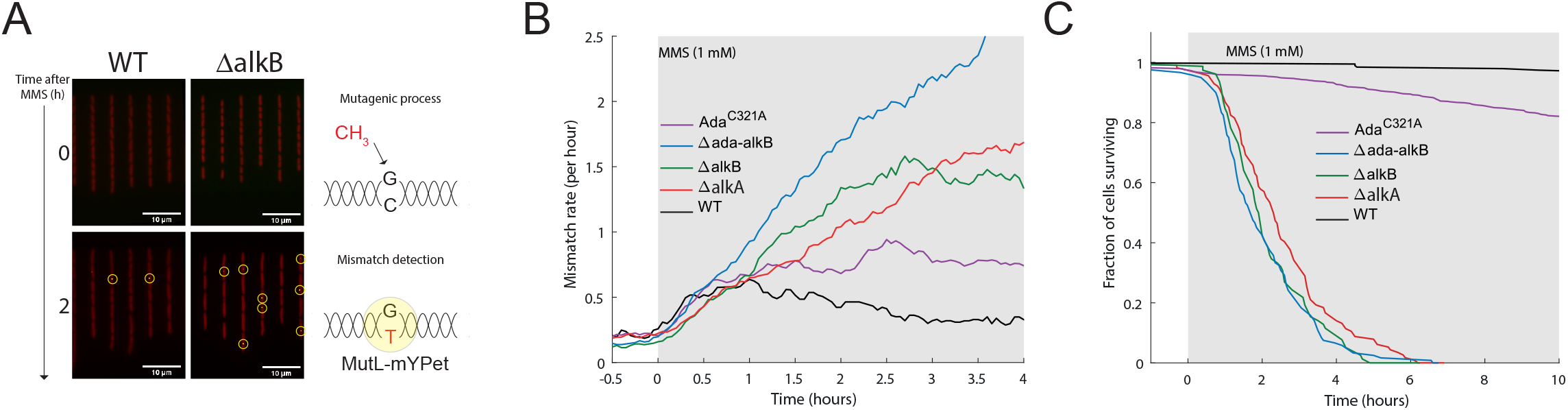
Contribution of Ada, AlkB and AlkA to the alkylating stress response. (A) Example of real-time imaging of DNA mismatch emergence. The addition of MMS in the fluidic system is mutagenic and results in nucleotide misincorporation during DNA replication. The DNA mismatch is recognised by the MutL-mYPet protein that forms fluorescent foci (yellow dots) and enables automated mismatch detection (yellow circles). Fluorescence of the segmentation marker mKate2 is shown in red. (B) Mismatch rate dynamics for strains *Δada-alkB* in blue (cells = 435), *ΔalkB* in green (cells = 347), *ΔalkA* in red (cells = 518), *ada*^*C321A*^ in purple (cells = 395), during constant 1 mM MMS treatment (shaded background) and compared with the WT strain shown in black (cells = 527). Mismatch rate curves have been smoothed using a moving average of 30 min. (C) Distribution of cell survival times during constant 1 mM MMS treatment for these same strains.

### Contributions of Ada, AlkB, and AlkA to cell survival

Although mutagenesis is essential for genome evolution, individual mutant cells that emerge during stress still need to survive in order to propagate their genetic innovations. We thus examined cell survival during MMS treatment. Despite the delay in the induction of alkylation repair, cell survival of the wild-type strain was essentially unaffected at 1 mM MMS (**Fig.4 C**), owing to the constitutively expressed DNA glycosylase Tag and DNA damage tolerance pathways that are controlled by the SOS response [39]. This was not the case for the *ΔalkB, ΔalkA*, and *Δada-alkB* deletion mutants, with less than 10% of cells surviving after 4 hours of constant MMS treatment for each of these strains (**Fig.4 C**). This result indicates that beyond a certain level of 3meA lesions (repaired by AlkA), and 3meC and 1meA lesions (repaired by AlkB), alternative repair and damage tolerance pathways cannot compensate for the lack of AlkA and AlkB. Failure to repair these lesions leads to DNA replication stalling [35,49,50], a process that is ultimately lethal to cells. On the other hand, 90% of *ada*^*C321A*^ cells were alive after 4 hours of constant MMS treatment, showing that Ada’s repair function protects predominantly against the mutagenic effects of alkylating stress rather than its toxicity.

### Cell-to-cell heterogeneity in the adaptive response leads to differences in genomic mutation rates

Our mismatch rates measurements imply that phenotypic heterogeneity in the DNA damage response causes cell-to-cell variation in mutation rates. Indeed, most DNA mismatches are repaired by the MMR system, but ~ 1% are overlooked and turn into stable mutations in the next round of replication [51]. However, whether differences in DNA mismatch rates truly reflect a genuine variation in mutation rates between cells remains unknown. To address this important point, we used fluorescence activated cell-sorting (FACS) to distinguish and collect cells that differentially activated the adaptive response after MMS exposure. We used a plasmid-based P_ada_-GFP reporter for the adaptive response that allowed us to identify two main subpopulations of cells after MMS treatment (**Supp.4**). One fluorescing, reflecting cells activating the response rapidly after MMS addition (ON) and one remaining non-fluorescent, reflecting cells with a delayed adaptive response (OFF) (**Supp.4**). We then sorted an identical number of 10^6^ cells from the two subpopulations and measured their respective mutation frequencies based on the number of colonies resistant to the antibiotic rifampicin (**Fig.5 A**). We found that the mutation frequency was significantly higher for the OFF than the ON subpopulations after 90 min of treatment with 1 mM (~ 1.5-fold difference), 3 mM (~ 5–fold difference) and 10 mM MMS (~4– fold difference) (**Fig.5 A**). Therefore, cell-to-cell variation in the timing of the adaptive response indeed causes substantial differences in genomic mutation rates. These results also confirm that the detection of MutL-mYPet foci as markers for DNA mismatches reports on the genomic mutation rates of single cells [10,39].

**Fig.5:**
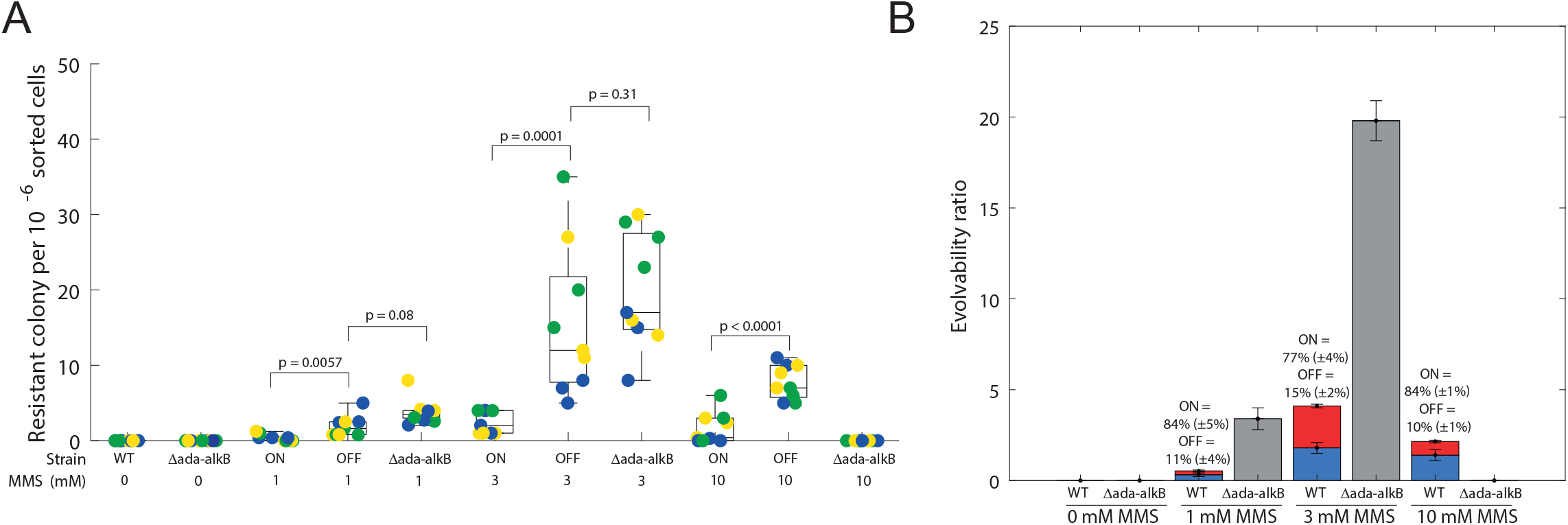
Consequences of the adaptive response heterogeneity on the mutation rate. (A) Boxplots showing the number of rifampicin resistant colonies for Ada-ON, Ada-OFF and *Δada-alkB* populations after 90 min treatment with different MMS concentrations (shown in mM). Each population was sorted according to defined sorting gates shown in supplementary.4. Biologically independent experiments (culture started from a distinct single colony) are grouped by colour. For each biological replicate, three rounds of sorting have been performed and plated on different rifampicin plates. P values were obtained with a two-tailed t-test. (B) Bar plot showing the evolvability ratio of each subpopulation after 90 min treatment with different MMS concentrations. The evolvability ratio has been defined as the product of the percentage of cells sorted in the total population and the rifampicin resistant mutant counts arising from this subpopulation. Averaged percentage of the subpopulation is shown for Ada-OFF and Ada-ON.

### Superior evolvability of the OFF subpopulation compared to Δada-alkB cells at high stress levels

To confirm that the P_ada_-GFP reporter activation was dependent on Ada, we also performed FACS with *Δada-alkB* cells. As expected, P_ada_-GFP remained inactivated independently of the MMS concentration (**Supp.4**). Furthermore, the mutation frequency of *Δada-alkB* cells was similar to that of the OFF subpopulation at 1 mM or 3 mM MMS. Therefore, wild-type cells that fail to activate the Ada response because of gene expression noise suffer the same mutagenic effects of alkylating stress as cells that lack the *ada* operon completely. However, *Δada-alkB* cells differed strongly from the OFF subpopulation at the higher dose of 10mM MMS (**Fig.5 A**). We did not detect any rifampicin-resistant colonies for the *Δada-alkB* strain after 10mM MMS treatment, whereas the OFF subpopulation generated a significant number of such colonies (**Fig.5 A**). We attribute the lack of mutant colonies to the extremely low survival of *Δada-alkB* cells in the presence of MMS. Thus, although *Δada-alkB* deletion promotes alkylation-induced mutagenesis (**Fig.4 B**), it also rapidly increases the likelihood of cell death (**Fig.4 C**). The disproportionate effect of the *Δada-alkB* deletion on the population dynamics therefore diminishes overall evolvability at high stress levels. Although OFF cells initially behave like *Δada-alkB* cells, they are capable of activating the Ada response eventually. This enables the repair of toxic DNA lesions that are otherwise lethal in *Δada-alkB* cells. The OFF subpopulation therefore accumulates mutations during the adaption delay but maintains chances of survival after response activation. These features make the OFF subpopulation a pool of increased genetic diversity.

### Evolvability as a trade-off between mutability and survival

We finally sought to address whether the OFF subpopulation contributes significantly to the evolvability of the whole population. This depends on several characteristics of the subpopulation, namely its size, mutation rate, and viability. By quantifying the number of rifampicin-resistant colonies relative to the abundances of the subpopulations, we found that the OFF subpopulation generates a substantial fraction of all viable mutants despite its small size (**Fig.5 B**). This analysis also demonstrated that evolvability is a trade-off between mutability and survival. The ON and OFF subpopulations both have a basal DNA damage tolerance owing to constitutively expressed DNA repair pathways and the SOS response. Increasing stress leads to higher mutation rates, but the lack of inducible DNA repair capacity in the OFF subpopulation means that cell survival drops disproportionately as the damage level rises. Because of this, the evolutionary benefit of the OFF subpopulation is maximal at intermediate damage level (3 mM MMS), where the subpopulation of 15% of cells generates 53% of the total rifampicin-resistant mutants.

## Discussion

Because emergence of mutations in a cell population is driven by rare and stochastic molecular events, mechanisms governing this process can be lost in the averaged result commonly gained with bulk experiments. We thus used a single-cell approach to study how regulatory dynamics of DNA repair genes influence mutation and cell survival, and ultimately impact evolvability of a cell population. Focusing on the adaptive response to DNA alkylation stress in *E. coli*, we found that, with the exception of AidB, the whole adaptive response regulon (i.e. Ada, AlkB, AlkA) is heterogeneously activated across isogenic cells during alkylating stress. Rather than a noisy genetic system, whereby gene expression heterogeneity is a side-effect of inaccurate control, the regulation of the adaptive response is orchestrated by a precise master regulator that divides an isogenic cell population into two defined subpopulations with distinct gene expression states. Interestingly, Ada levels are upregulated thousandfold in response to alkylation damage, yet cells expressing the non-functional Ada^C321A^ mutant that is defective in O^6^meG lesion repair are not sensitised to alkylation damage and exhibit only slightly increased mismatch rates. The benefit of high Ada numbers could be an increased robustness to gene expression noise after response activation. Indeed, we found that fluctuations in Ada expression are accurately propagated to AlkA. Conversely, the expression of all adaptive response genes is very low before the response induction. The low basal level of Ada combined with positive feedback amplification in the presence of alkylation stress creates a stochastic switch where the infrequent expression of a single Ada molecule is the trigger that turns cells from the OFF to the ON state. The very low basal abundance of AlkA and AlkB was unexpected in light of their delayed induction and importance for cell survival. This reinforced the view that a lack of DNA alkylation repair capacity in a subpopulation of cells serves a particular purpose that provides a greater benefit than its cost to instantaneous cell fitness.

We show that heterogeneity in the adaptive response represents the phenomenon of stress-induced mutagenesis, whereby cells poorly adapted to their environment increase their mutation rates. Whether this is an evolvability strategy *per se* or an unavoidable consequence of the selection for survival has been brought into question [17,52]. Nonetheless, the functional benefit of stress-induced mutagenesis relies on the ability of cells to propagate any mutations that are generated during stress. Indeed, alternative DNA repair and damage tolerance mechanisms, such as constitutively-expressed DNA glycosylases and the translesion synthesis DNA polymerases of the SOS response can rescue early failures to repair toxic alkylation lesions [39]. However, when replication-stalling lesions saturate alternative repair strategies, the adaptive response becomes necessary for survival [39]. Our study demonstrated that cells with a delayed adaptive response have an elevated rate of nascent mutations and maintain the capacity to propagate these mutations if they eventually activate the adaptive response. In this way, the transient hypermutable subpopulation generated by stochastic regulation of DNA alkylation repair increases the evolvability of the whole population.

## Acknowledgments

The authors thank members of the Uphoff group and the group of David Sherratt for insightful discussions. M.S.V would like to thank Nicolas Flaugnatti, Pierre Santucci and Yassine Cherrak for their helpful comments on this work. Research in the Uphoff laboratory is funded by a Sir Henry Dale fellowship (206159/Z/17/Z) and a Wellcome-Beit prize (206159/Z/17/B) by the Wellcome Trust, and a Research Prize Fellowship of the Lister Institute of Preventative Medicine. S.U. holds a Hugh Price Fellowship at Jesus College, Oxford. M.S.V. is funded by a Human Frontiers Science Programme long-term fellowship (LT000092/2019-L) and holds an EMBO non-stipendiary long-term fellowship (ALTF 1035-2018).

## Materials and Methods

### Construction of strains and plasmids

All strains were derived from *Escherichia coli* K12 AB1157. C-terminal msCFP3, mYPet and HaloTag fusions were inserted with a flexible 11 amino acids linker (SAGSAAGSGEF) at the endogenous chromosomal loci by λ-red recombination using plasmids pSU003 [37], pRod50 [53] and pSU005 [54]. The λ-red insertions are flanked by Flp site-specific recombination sites (frt) that allow removing the antibiotic resistance gene using Flp recombinase from plasmid pCP20[55]. After recombination, all λ-red insertions were confirmed by colony PCR and the alleles were moved into new strains by P1 phage transduction. The dual reporter strains carrying the *P*_*ada*_*-CFP* reporter has been described in [37]. It is a transcriptional fusion made of a single copy of the *ada* promoter followed by the CFP fast-maturing variant SCFP3A inserted at the chromosomal *intS* site (~150 kb downstream from the native *ada* gene). The *ΔalkB* deletion strain was obtained from the Coli Genetics Stock Center (CGSC 9779) and moved into other strains by P1 phage transduction. The *ΔalkA* and *Δada-alkB* mutants were engineered by λ-red recombination. The chromosomal *ada*^*C321A*^ point mutant has been engineered by λ-red recombination using plasmid pMV010. This plasmid is derived from plasmid pMV001 that has been synthesized with GeneArt Gene Synthesis (ThermoFisher Scientific). pMV001 carries the *P*_*ada*_*-ada-alkB* operon where codons encoding Ada C38 and C321 have been replaced to encode A38 and A321. pMV010 underwent site-directed mutagenesis (NEBaseChanger) to restore the original codon encoding C38. An additional chloramphenicol resistance cassette was inserted downstream *alkB* into pMV010 to select for recombinant cells after λ-red recombination into *Δada-alkB*.

### Cell culture

Strains were streaked from frozen glycerol stocks onto LB agarose with appropriate antibiotic selection. A single colony was used to inoculate LB and grown for 6-7 hours. The cultures were then diluted 1:1000 into supplemented M9 minimal medium containing M9 salts (15 g/L KH2PO4, 64 g/L Na2HPO4, 2.5 g/L NaCl, 5.0 g/L NH4Cl), 2 mM MgSO4, 0.1 mM CaCl2, 0.5 µg/ml thiamine, MEM amino acids, 0.1 mg/ml L-proline, 0.2% glucose. Cultures were grown overnight to stationary phase, then diluted 1:50 into supplemented M9 medium and grown to OD600= 0.2.

### Single-molecule counting microscopy

Cells were treated as described in the cell culture section until OD600 = 0.2 and resuspended into 100 μl of supplemented M9 minimal medium. Cells expressing HaloTag fusions were labelled with TMR ligand (Promega) following the procedure previously described in [54]. Briefly, 5 μl of 2.5 μM TMR ligand was added to the cell resuspension and incubated for 30 min at 25°C. TMR dyes were then removed with four rounds of washing. Cells were resuspended into 1 ml of supplemented M9 minimal medium and incubated at 37°C for 30 minutes. In order to stop protein diffusion, cells were pelleted and resuspended into 2.5% paraformaldehyde in PBS buffer and fixed for 30 min at room temperature. Fixed cells were centrifuged, concentrated 10-fold and 1 μl of the cell resuspension was spotted on an agarose pad. Single-molecule imaging was performed using a custom-built total internal reflection fluorescence (TIRF) microscope under oblique illumination at room temperature. Epifluorescence illumination was used to ensure that Halo-tagged proteins were detected within the whole cell. To reduce both background noise and the probability of missed detection due to TMR stochastic blinking we used exposure time of 1 sec and acquired 5 frames per sample under continuous 561 nm excitation at 0.2 kW.cm^-2^. TMR-labelled Halo-tagged proteins are initially in the fluorescent state [54], hence to avoid loss of detection of fast bleaching TMR we set up our camera with TTL mode, such that the first frame acquired contains all fluorescent molecules. The 5 frames were then averaged in a single image using the Z-projection function of ImageJ [56] (Projection type: averaged intensity) and fluorescent spots were counted manually.

### Single-cell microfluidic-based microscopy

Cells were treated as described in the cell culture section until OD600 = 0.2. 0.85 mg/mL of surfactant pluronic F127 (Sigma Aldrich) was added to the culture to avoid cell aggregation in the microfluidic device. The microfluidic single-cell imaging device was previously designed and experiments were performed as described in [39]. In addition to fluorescent reporters of the adaptive response, strains used for single-cell measurement constitutively expressed the fluorescent protein mKate2 and carried an *flhD* gene deletion to remove flagellum motility. Imaging was performed on a Nikon Ti Eclipse inverted fluorescence microscope equipped with perfect focus system, 100x NA1.45 oil immersion objective, sCMOS camera (Hamamatsu Orca Flash 4), motorized stage, and 37°C temperature chamber (Okolabs). Fluorescence images were automatically collected using NIS-Elements software (Nikon) and an LED excitation source (Lumencor SpectraX). Time-lapse movies were recorded at 3-min intervals with exposures time of 75ms for msCFP3, 100ms for mKate2 and 300ms for mYPet, using 50% LED excitation intensities.

### Data analysis

Microscopy movies were analyzed using custom MATLAB software to segment cells based on cytoplasmic mKate2 fluorescence. Only mother cells at the end of each channel were included in the analysis. Cell deaths were manually detected when growth ceased, or when time traces terminated abruptly because cell filamentation led to the disappearance of the cell from the growth channel. mYPet and CFP reporters intensities were calculated from the average pixel intensities inside the segmented cell area and subtracting the background signal outside of cells. Detection of MutL-mYPet foci for mismatch rate determination was performed with a spot-finding algorithm [57]. When foci persisted for several frames, only the first frame was counted as a mismatch event. Mismatch rates were calculated by dividing the number of observed mismatch events by the observation time interval. Cell-average time traces of mismatch rates were generated by dividing the number of mismatch events by the number of observed cells in each frame. Pearson correlation coefficients were calculated using the MATLAB corrcoef function. Cross-correlations between fluorescence signals were calculated using the MATLAB xcorr function.

### FACS and rifampicin assays

4ml of M9 supplemented with kanamycin (25ug/ml) was inoculated with a single colony of WT (SU828) or Δada-alkB (SU829). SU828 and SU829 contain a constitutive mKate2 segmentation marker and a kanamycin resistant plasmid (pUA139) encoding a P_ada_-GFP reporter (obtained from the Uri Alon’s reporter library [58]). At OD600 = 0.2, cells were treated with 1mM, 3mM or 10mM MMS for 90 min. 1 ml of cells were then washed two times by centrifugation and resuspended into 1X PBS to remove residual MMS. Cells were diluted into 5 ml 1X PBS and sorted and analysed with a S3e(tm) Cell Sorter and ProSort(tm) Software (BioRad). Fluorescence intensities of cells were measured using 488 nm and 561 nm lasers. Signals were collected using the emission filters FL1 (525/30 nm) and FL3 (615/25 nm) for GFP and mKate2, respectively. Voltages of the photomultipliers were 500, 300, 600, and 720 volts for FSC (forward scatter), SSC (side scatter), FL1, and FL3, respectively. Histograms obtained (Cells count versus GFP fluorescence) were gated on the population of interest and cells were sorted in 5ml 1X PBS at a rate of 10000 particles per second. Cells were diluted into 10 ml of LB and incubated at 37C for one hour. Cells were centrifuged (4500 rpm for 10 min) and resuspended into 1ml of LB before being plated on freshly prepared LB agar plates with 20ug/ml rifampicin. After over night incubation at 37C, the number of rifampicin resistant clones was counted and divided by the number of cells sorted to define the rifampicin resistance frequency. Measurements were carried out 3 times with bacteria from different plates.

## Figure Legends

**Supp.1:**
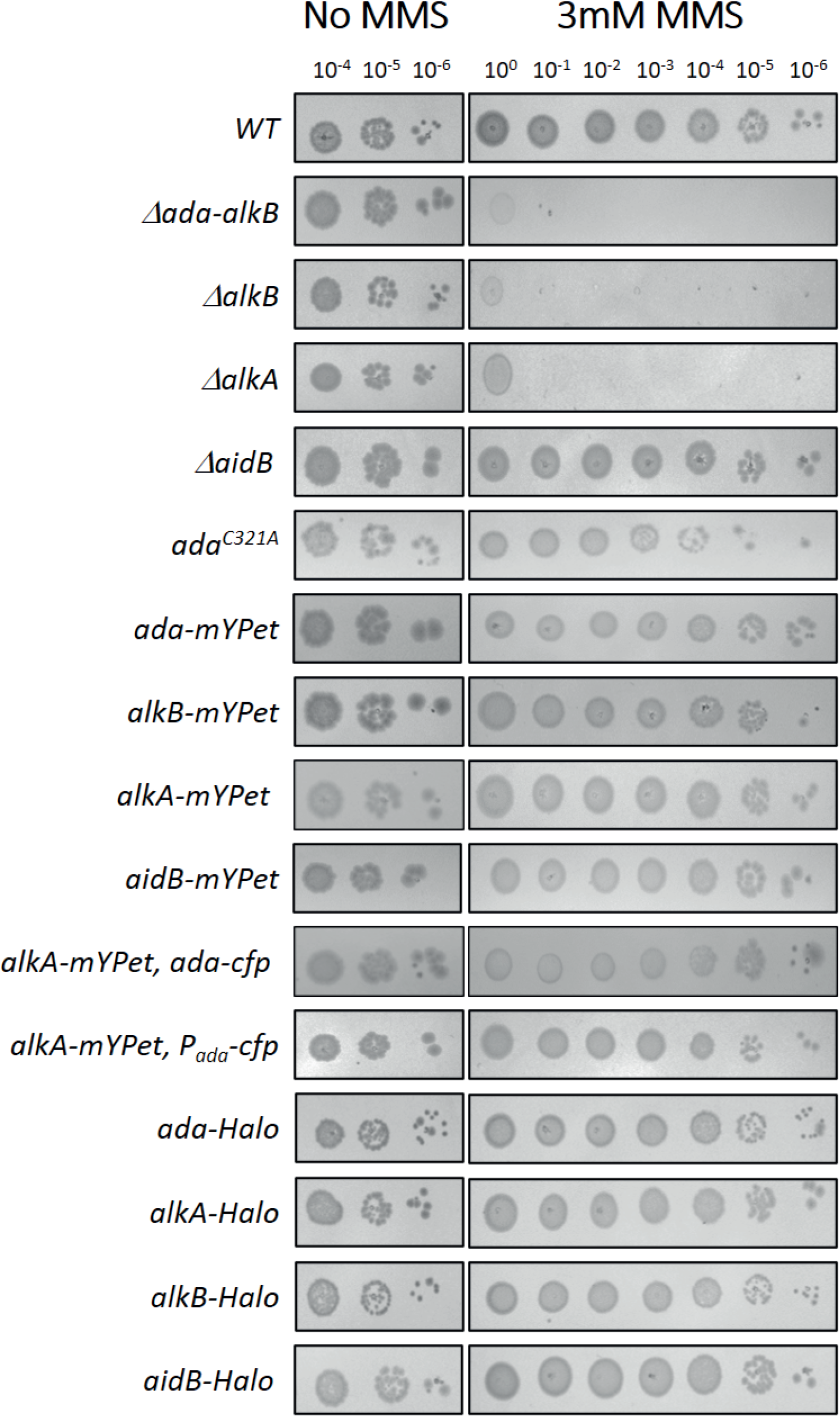
MMS sensitivity assays. To assess the functionality of translational and transcriptional reporters used in this study, we performed MMS sensitivity tests by spotting 10-fold serial dilutions of over-night cell cultures on LB and LB + 3mM MMS plates and compared growths after over-night incubation at 37C. Note that the P_ada_-cfp reporter refers to an ectopic version of the ada promoter, the native *P*_*ada*_*-ada-alkB* locus is unaltered in this strain.

**Supp.2:**
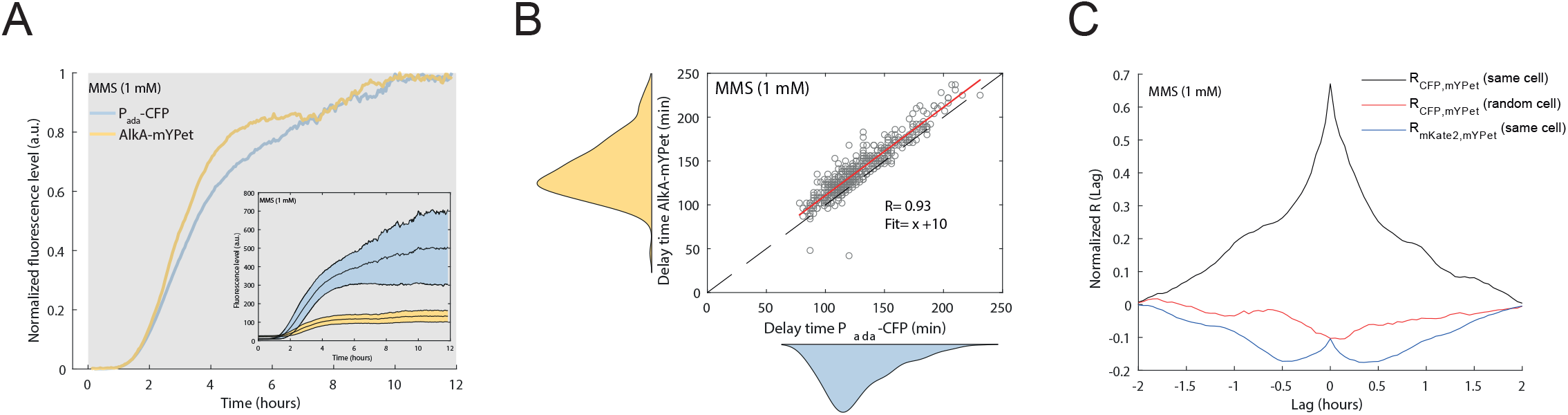
Dual reporter assays of P_ada_-CFP and AlkA-mYPet activation upon 1 mM MMS treatment. (A) Fluorescence of each single-cell (cells = 401) has been averaged, normalized and subtracted from their level at time = 0h (addition of MMS in the fluidic system). Inset shows the original signals and their standard deviations about the mean (coloured regions). (B) Correlation plot showing delay times between 1.5 mM MMS addition and response activation for P_ada_-CFP and AlkA-mYPet. Each circle represents one cell. R is the Pearson coefficient. The red line shows the best linear fit. (C) Cross-correlations of P_ada_-CFP and AlkA-mYPet signals between 9 and 11 hours after 1.5 mM MMS addition. The average of each individual cross-correlation between the mYPet and CFP signals from the same cell is represented by the black curve, whereas the red curve represents the average from two random cells and indicates that the correlation between Ada and AlkA is specific of their respective cell. The blue curve represents the average of each individual cross-correlation between the AlkA-mYPet signal and the segmentation marker mKate2 signal from the same cell and indicates that the correlation is independent of the fluorescence fluctuations due to cell elongation during the cell cycle.

**Supp.3:**
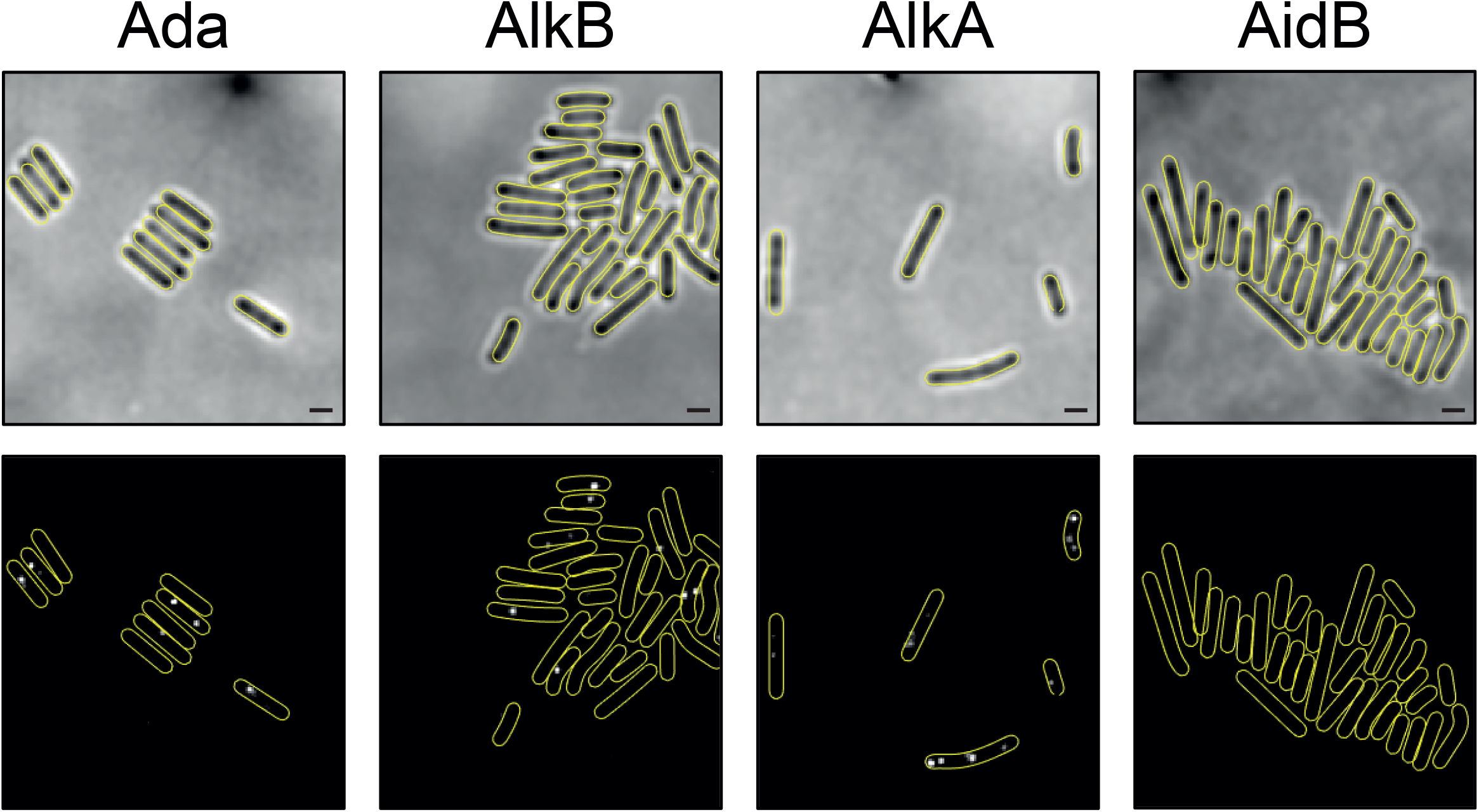
Basal level of the adaptive response proteins. Example of single molecule spots detected within chemically fixed cells after in vivo HaloTag labelling with TMR ligand. Upper panel = brightfield, lower panel = 561 nm channel (averaged stacks). Scale bar = 1μm.

**Supp.4:**
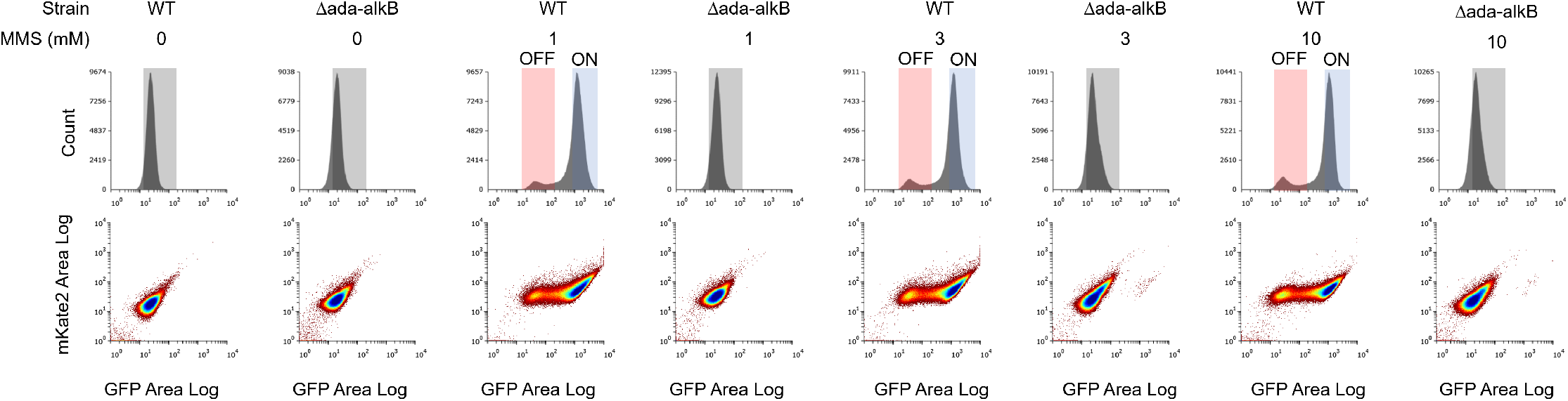
Detection and sorting of Ada-ON and Ada-OFF subpopulations. Flow-cytometry was performed on the WT and *Δada-alkB* strains, both carrying a plasmid-based P_ada_-GFP reporter and a segmentation marker mKate2. The segmentation marker enables to exclude debris, dead cells and contaminants from the analysis. In absence of MMS treatment, WT cells exhibit a unimodal distribution that is used to define the inactivated-population gate (OFF). After 90 min treatment with 1, 3 or 10 mM MMS, WT cells exhibit a bimodal distribution reflecting the subpopulation of cells delaying (gate OFF) or activating (gate ON) the adaptive response. The *Δada-alkB* remains inactivated independently of the MMS concentration and allows us to control that the P_ada_-GFP reporter is dependent on the Ada production.

**Table S1:**
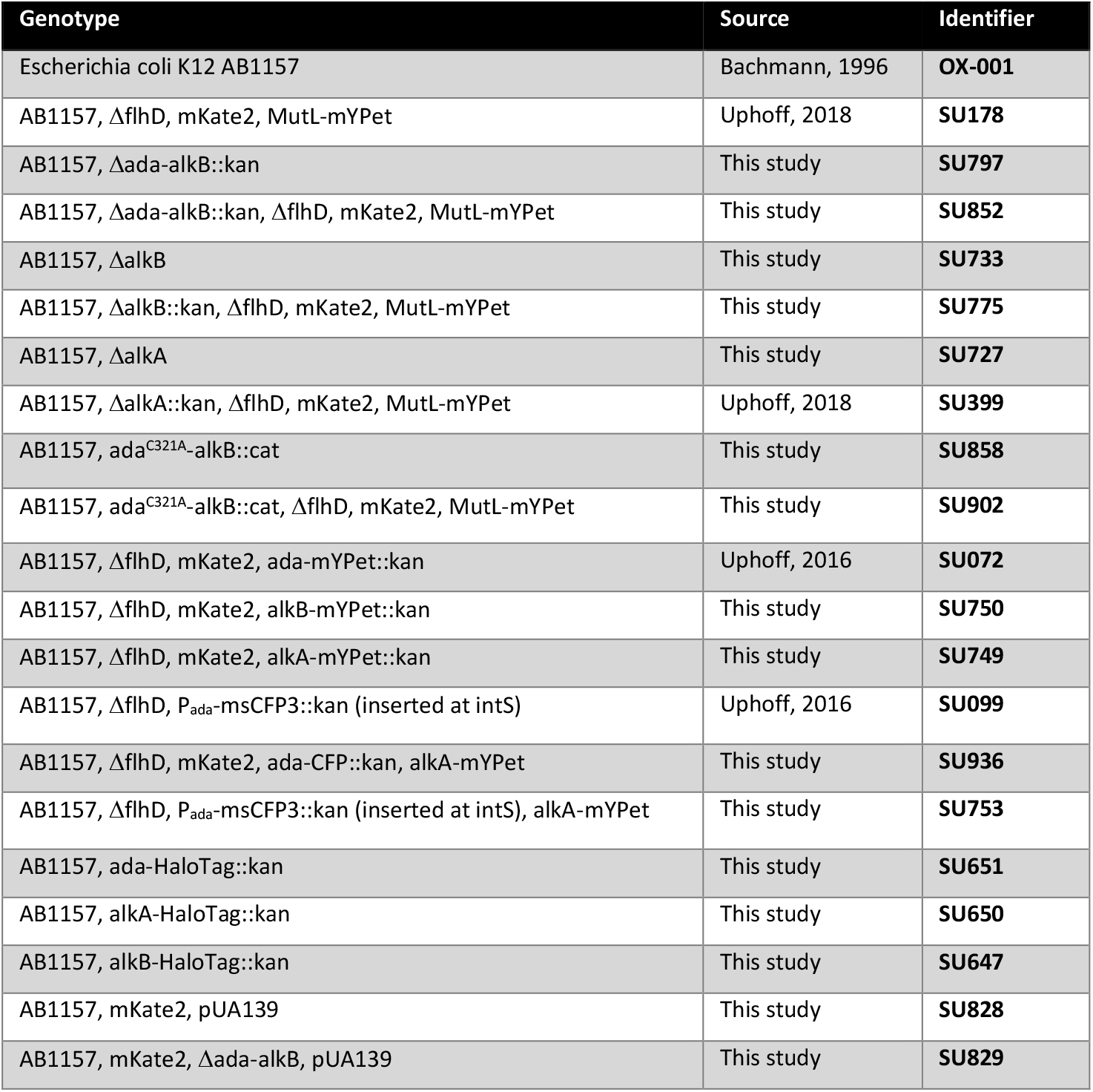
Strains used in this study.

| Genotype | Source | Identifier |
| --- | --- | --- |
| Escherichia coli K12 AB1157 | Bachmann, 1996 | **OX-001** |
| AB1157, ΔflhD, mKate2, MutL-mYPet | Uphoff, 2018 | **SU178** |
| AB1157, Δada-alkB::kan | This study | **SU797** |
| AB1157, Δada-alkB::kan, ΔflhD, mKate2, MutL-mYPet | This study | **SU852** |
| AB1157, ΔalkB | This study | **SU733** |
| AB1157, ΔalkB::kan, ΔflhD, mKate2, MutL-mYPet | This study | **SU775** |
| AB1157, ΔalkA | This study | **SU727** |
| AB1157, ΔalkA::kan, ΔflhD, mKate2, MutL-mYPet | Uphoff, 2018 | **SU399** |
| AB1157, ada<sup>C321A</sup>-alkB::cat | This study | **SU858** |
| AB1157, ada<sup>C321A</sup>-alkB::cat, ΔflhD, mKate2, MutL-mYPet | This study | **SU902** |
| AB1157, ΔflhD, mKate2, ada-mYPet::kan | Uphoff, 2016 | **SU072** |
| AB1157, ΔflhD, mKate2, alkB-mYPet::kan | This study | **SU750** |
| AB1157, ΔflhD, mKate2, alkA-mYPet::kan | This study | **SU749** |
| AB1157, ΔflhD, P<sub>ada</sub>-msCFP3::kan (inserted at intS) | Uphoff, 2016 | **SU099** |
| AB1157, ΔflhD, mKate2, ada-CFP::kan, alkA-mYPet | This study | **SU936** |
| AB1157, ΔflhD, P<sub>ada</sub>-msCFP3::kan (inserted at intS), alkA-mYPet | This study | **SU753** |
| AB1157, ada-HaloTag::kan | This study | **SU651** |
| AB1157, alkA-HaloTag::kan | This study | **SU650** |
| AB1157, alkB-HaloTag::kan | This study | **SU647** |
| AB1157, mKate2, pUA139 | This study | **SU828** |
| AB1157, mKate2, Δada-alkB, pUA139 | This study | **SU829** |

**Table S2:**
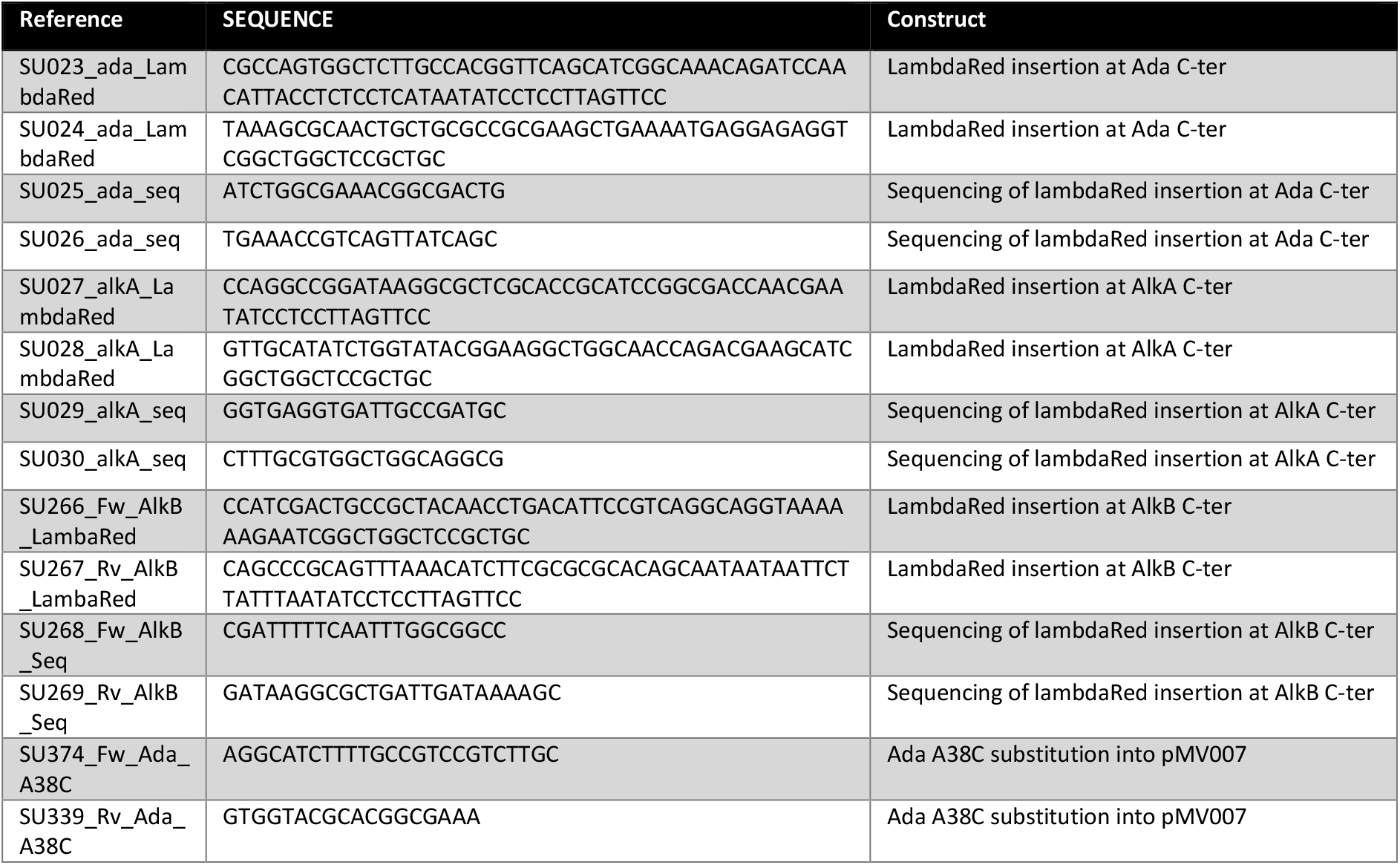
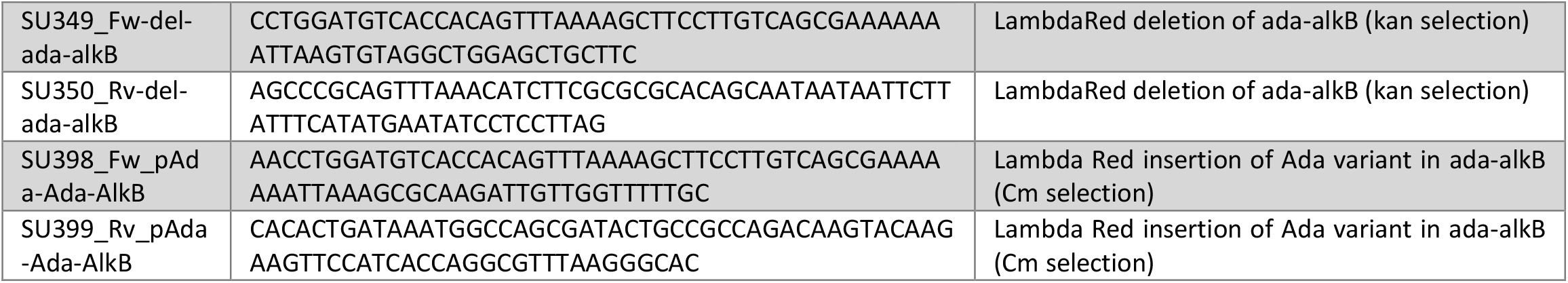
Primers used in this study.

## Notes

### Competing Interest Statement

The authors have declared no competing interest.

